# *SIRT2, TFEB, CNIH3, ADRM1*, AND *TM9SF4* AS POTENTIAL BIOMARKERS OF RESISTANCE TO CRIZOTINIB AND POOR PROGNOSIS IN LUNG ADENOCARCINOMA: IMPLICATIONS FOR TARGETED THERAPY AND IMMUNOTHERAPY

**DOI:** 10.1101/2025.07.10.662953

**Authors:** Édgar Villena Maciá, Maissaa Bouchakour-Bouchaqor, Paula Sánchez-Olivares, Fernando Andrés-Pretel, Eva M. Galán-Moya

## Abstract

**Background:** Lung adenocarcinoma (LUAD) with ALK mutations benefits from targeted treatment with ALK tyrosine kinase inhibitors, such as crizotinib. This therapy has been shown to be more effective than traditional chemotherapy, with improved tolerance in patients. However, resistance to crizotinib can develop, limiting its long-term efficacy. The aim of this study is to identify deregulated genes in crizotinib-resistant ALK-mutated lung adenocarcinoma cell lines. This will allow us to select patients who are more likely to benefit from crizotinib and help design new therapeutic strategies to overcome resistance, ultimately leading to better clinical outcomes.

**Methods:** An *in silico* study using genomic data from crizotinib-sensitive and crizotinib-resistant LUAD cells was conducted. Through statistical analysis, genes with differential expression in crizotinib resistant cells were identified. These genes were classified by their biological function using Enrichr, and those with the highest amplification frequency according to cBIOPORTAL were selected. Next, gene expression association with prognosis and with immune cell infiltration were investigated using Kaplan-Meier plotter and TIMER databases, respectively. Gene expression analysis in normal and tumoral tissues was explored using TNM plot and potential therapeutic agents against the identified targets were identified from several databases.

**Results:** We found 9 genes overexpressed in resistant LUAD cells, with statistically significant cell function, to be amplified and associated with poor prognosis. Among them, SIRT2, TFEB, TMPSF4, CINH3 and ADRM1 display an immunosuppressed TME and TMPSF4, CINH3 and ADRM1 are overexpressed in tumoral tissue. Some of them, such as SIRT2, have therapeutic agents already approved in other context (gefitinib in EFGR mutated lung cancer).

We identified nine genes that are overexpressed in crizotinib-resistant cell lines and are associated with poor prognosis. Additionally, we observed differences in immune infiltration patterns based on the specific overexpressed gene. These findings suggest potential avenues for therapeutic intervention, such as the use of combination therapies with crizotinib, including immunotherapy or targeted therapies aimed at resistance pathways. These strategies could enhance treatment efficacy in tumors expressing these genes.

**Conclusions:** *SIRT2, TFEB, TM9SF4, CINH3,* and *ADRM1* are potential biomarkers for crizotinib-resistant and worse prognosis in LUAD. Our findings provide valuable insights into the mechanisms of crizotinib resistance and potential avenues for therapeutic intervention.

**Highlights:**

- *In silico* analysis identified nine overexpressed genes in crizotinib-resistant lung adenocarcinoma cell lines, all associated with worse prognosis.
- Tumor levels of *SIRT2, TFEB, TM9SF4, CINH3,* and *ADRM1* correlate with higher infiltration of immunosuppressive populations and, the last three, with lower infiltration of immune-promoting cells.
- These genes are potentially druggable targets in the context of crizotinib resistance, with existing treatments that could be explored.
- The tumor microenvironment (TME) may play a significant role in these tumors, suggesting that immunotherapy could be a viable treatment option in this setting.

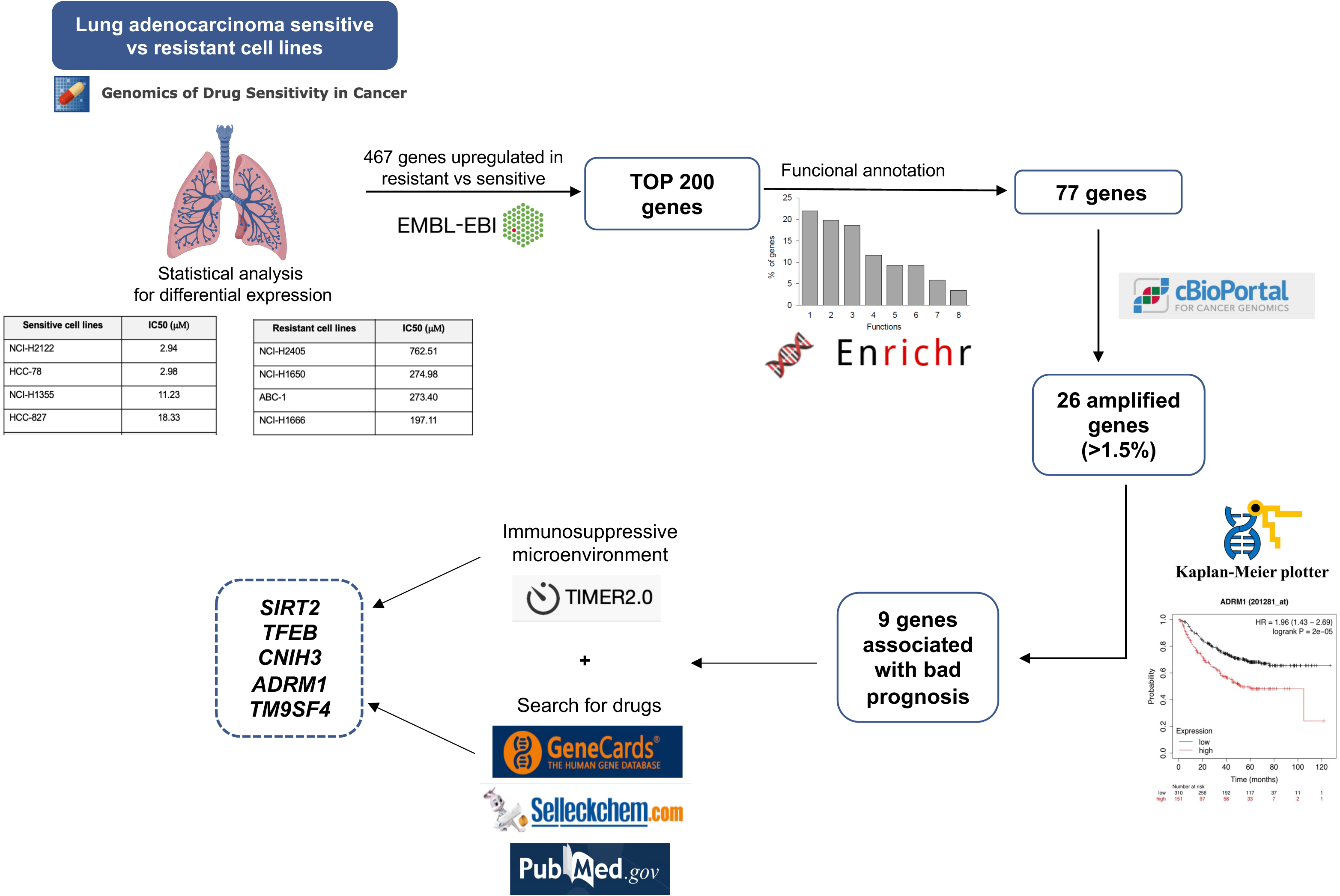

## 1. Introduction

Lung cancer is the leading cause of cancer death and the second cause of cancer incidence worldwide. Up to 90% of new cases are related to tobacco use, which plays a role in tumorigenesis and the development of metastases. The rest of the cases are related to exposure to radon, human immunodeficiency virus, tuberculosis, etc. There is also a certain degree of genetic susceptibility, with first-degree relatives presenting up to four times the risk of suffering it, regardless of smoking habits [1].

Non-microcytic histology is present in 80-85% of these tumors, including adenocarcinoma (LUAD), squamous and large cell. Adenocarcinoma is associated with translocations of the *ALK* gene (anaplastic lymphoma kinase) in 3-5% of cases and is characterized by a high risk of metastasis to the central nervous system, with more than 30% presenting brain metastases at diagnosis, and 50% developing them [1, 2]. This gene is located on chromosome 2 and encodes a receptor tyrosin kinase of the insulin family, which is expressed in central and peripheral nervous tissue during embryogenesis The formation of a fusion protein through a cromosomal translocation is the most common mechanism of *ALK* kinase activation and overexpression. The translocation of the *ALK* gene with the *EML4* gene (echinoderm microtubule-like protein 4) results in the formation of the *EML4*-*ALK*. This fusion gene, in locus 2p21, encode proteins that activate intracellular signaling pathways, such as MAPK (Mitogen-Activated Protein Kinase), JAK-STAT (Janus Kinase - Signal transducer and activator of transcription proteins) or PI3K (Phosphoinositide 3-kinase) and, therefore, favor tumor proliferation and survival. This translocation is more frequent in LUAD with a cribriform pattern or with signet ring cells and usually appears in young patients, with little or no history of smoking and at an advanced stage [3,4,5,6].

Within the therapeutic arsenal against *ALK*, first generation (crizotinib), second generation (ceritinib, alectinib, brigatinib), and third generation (lorlatinib) tyrosin kinase inhibitors (TKIs) are available. The first drug approved against *ALK* was crizotinib in 2011 by the FDA (Food and Drug Administration) and in 2015 by the EMA (European Medicines Agency). This approval was based on the results of the phase III PROFILE 1014 trial, which demonstrated better PFS (progression free survival) compared to platinum-based chemotherapy [4]. Regarding the response at the central nervous system level, crizotinib also demonstrated better response compared to chemotherapy [5].

However, crizotinib also has characteristic adverse effects, such as visual disturbances (up to 60% depending on the series), hepatic disturbances (up to 40%), peripheral edema (up to 30%), and QT segment lengthening or bradycardia [6].

At the research level, in second line and post progression after crizotinib, cells develop resistance mutations, which can be on target, which are those that induce amplifications of the *ALK* gene, and off target, related to upregulation and bypass signaling pathways. For example, patients with EML4-*ALK* variant 3a/b, who progress to crizotinib more frequently, present mutations in G1202R of *ALK*, which induces resistance to 2nd generation inhibitors, so they should be treated with 3rd generation inhibitors, such as lorlatinib [7].

In this research, we will try to identify crizotinib resistance genes in *ALK*-mutated lung adenocarcinoma, in order to identify those patients who do not benefit from first-line treatment with crizotinib monotherapy and to discover strategies to overcome this resistance.

## 2. Materials and methods

### 2.1. Selection of crizotinib sensitive and resistant lung adenocarcinoma cell lines

LUAD cell lines resistant and sensitive to crizotinib were searched, to subsequently analyze differences in gene expression between the two groups. To do so, we searched for cell lines treated with crizotinib in the Genomic of Drug Sensitivity in Cancer database (https://www.cancerrxgene.org), specifically LUAD. We then sorted them according to their IC50, and selected the cell lines most sensitive and resistant to crizotinib, specifically the four most sensitive and the four most resistant.

### 2.2. Differential gene expression between crizotinib sensitive and resistant cell lines

We downloaded the gene expression of all eight cell lines from the EMBL-EBI database to find gene expression differences between the four most resistant and four most sensitive cell lines. Using statistical methods, we have analyzed the genes showing statistically significant differences between the resistant and sensitive cell lines.

### 2.3. Functional analysis in lung adenocarcinoma cell lines resistant to crizotinib

The functional analysis of the genes that show differential expression in resistant versus sensitive lines was performed in order to analyze the cellular functions related to treatment resistance. It was done using Enrichr and we have selected the genes that are related to statistically significant cellular functions.

### 2.4. Identification of amplified genes in patients with crizotinib-resistant lung adenocarcinoma and differential analysis with healthy tissue

Once we have selected the genes with a statistically significant function in relation to crizotinib resistance, we have performed a data analysis in patients with *ALK*-mutated lung adenocarcinoma using the cBIOPORTAL platform, where amplification and mutation percentages in tumor tissue are analyzed. We have selected those with an amplification percentage higher than 1.5%. In addition, of these genes, we have performed a differential expression analysis between tumor and non-tumor tissue, using the TNMplot platform, due to a potential higher toxicity of crizotinib in patients with non-tumor tissue overexpressing these genes.

### 2.5. Prognostic analysis of overexpressed genes in crizotinib-resistant lung adenocarcinoma

To assess the significance of the overexpression of these genes in patients with crizotinib-resistant LUAD, we used the Kapplan-Meier plotter platform, which relates these genes to prognostic data [11]. Specifically, we have classified them according to their prognosis in terms of overall survival (OS) and PFS.

### 2.6. Association with immune infiltration

Once the genes associated with worse prognosis were identified, their correlation with immune infiltrates was evaluated using the TIMER2.0 platform. The immune populations analyzed in the TME were effector T cells (CD8), regulatory T cells (T-regs), macrophages, neutrophils and dendritic cells (DC) using the expression profiles by EPIC, TIMER, or CIBERSORT.

### 2.7. Gene-targeted drugs in crizotinib resistant cell lines

Of the genes overexpressed in crizotinib-resistant cell lines and which are associated with worse prognosis, we have searched for treatments that modulate the action of these genes, both those already approved for other indications and those under investigation. To do so, we accessed different databases: gene card, selleck chem, clinical trial and pubmed. We searched for treatments used in the clinic and in the experimental phase that regulate the action of overexpressed genes in crizotinib-resistant cell lines, which are associated with a worse prognosis.

## 3. Results

### 3.1. Functional analysis in lung adenocarcinoma cell lines resistant to crizotinib

To search for genes differentially expressed in cells sensitive and resistant to crizotinib, we searched for cell lines of LUAD treated with crizotinib in the Genomic of Drug Sensitivity in cancer database. The 47 cell lines found were sorted according to their IC50 to crizotinib, and the four most sensitive and the four most resistant were chosen to pursue the study (Table 1). For these eight cell lines, gene expression data was downloaded from EMBL-EBI database and genes with the highest expression in the resistant cell lines were identified. A total of 467 genes were found to be differentially overexpressed in the resistant cell lines and then, the 200 with the highest differential expression were selected (Supplementary Table 1).

**Table 1.** Crizotinib sensitivity of lung adenocarcinoma cell lines. The four most sensitive and the four most resistant cell lines are shown, according to data from the Genomics of Drug Sensitivity in Cancer database.

| Sensitive cell lines | IC50 (μM) |
| --- | --- |
| NCI-H2122 | 2.94 |
| HCC-78 | 2.98 |
| NCI-H1355 | 11.23 |
| HCC-827 | 18.33 |

| Resistant cell lines | IC50 (μM) |
| --- | --- |
| NCI-H2405 | 762.51 |
| NCI-H1650 | 274.98 |
| ABC-1 | 273.40 |
| NCI-H1666 | 197.11 |

| Sensitive cell lines | IC50 (μM) |
| --- | --- |
| NCI-H2122 | 2.94 |
| HCC-78 | 2.98 |
| NCI-H1355 | 11.23 |
| HCC-827 | 18.33 |
| Resistant cell lines | IC50 (μM) |
| NCI-H2405 | 762.51 |
| NCI-H1650 | 274.98 |
| ABC-1 | 273.40 |
| NCI-H1666 | 197.11 |

To identify altered biological functions in crizotinib resistant cell lines, these 200 genes were used for functional annotation analysis using Enrichr database. Eight biological functions were identified as the most altered, accounting for 77 genes in total (Fig.1). The most represented altered function was regulation of metabolic processes. This function had a *p-value* of 0.00207659 and includes up to 22.1% of the total number of genes (19 genes). The second most represented function (19.8%, 17 genes) was regulation of transcriptional and translational processes (*p-value*= 0.00148042), followed by protein post-translational modification (*p-value* 0.000178), with 18.7% of the total of genes (16 genes).

**Figure 1.**
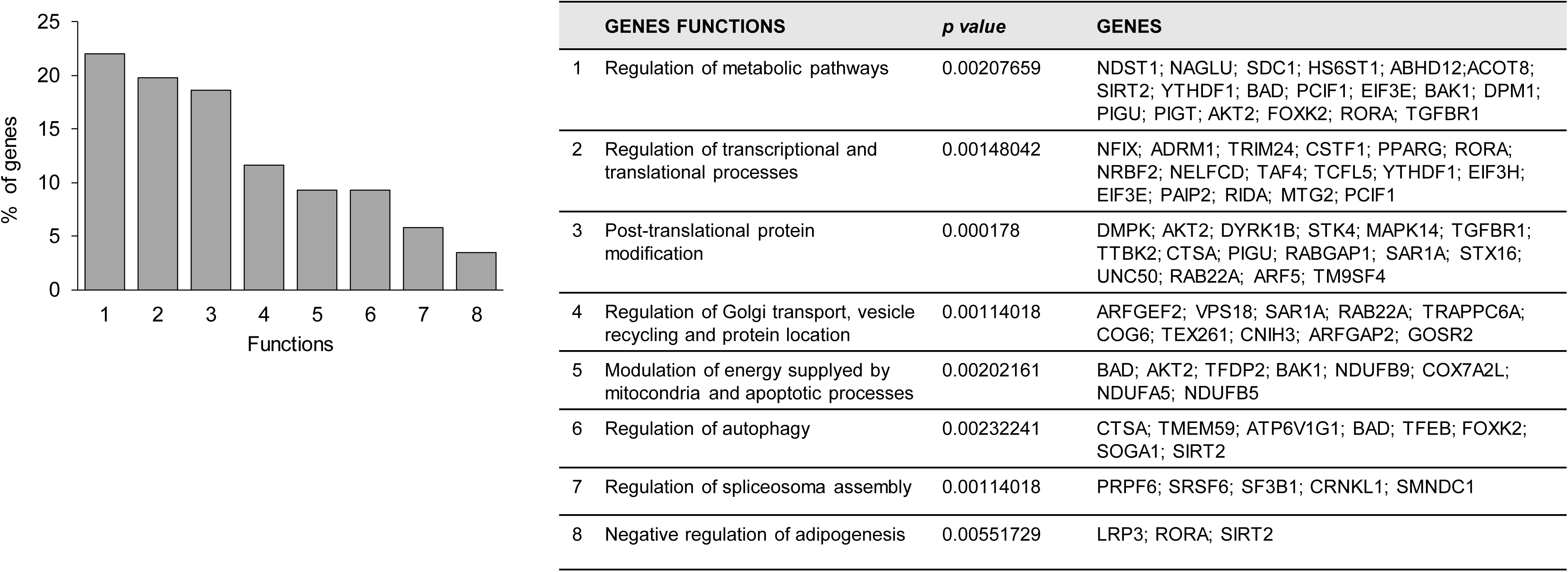
Functional annotation analysis of the 200 most upregulated genes in crizotinib-resistant lung adenocarcinoma cell lines. Statistically significant functional categories were selected based on *p-value*. Functional categories: 1. Post-translational protein modifications, 2. Regulation of spliceosome assembly, 3. Regulation of Golgi transport, vesicle recycling and protein localization, 4. Regulation of transcriptional and translational processes, 5. Modulation of energy supplied by mitochondria and apoptotic processes, 6. Regulation of metabolic pathways, 7. Regulation of autophagy, 8. Negative regulation of adipogenesis.

Regulation of transport mechanisms at the level of the Golgi apparatus, vesicular recycling and protein localization accounted for 11.62% (10 genes, *p-value* 0.00114018), modulation of energetic processes derived from mitochondrial metabolism and apoptosis, 9.3% (8 genes, *p-value* 0.00202161), autophagy, 9.3% (8 genes, *p-value* 0.00232241), spliceosome assembly, 5.81% (5 genes, *p-value* 0.00114018) and negative regulation of adipogenesis, 3.37% (3 genes, *p-value* 0.00551729).

### 3.2. Prognostic analysis of genes overexpressed in crizotinib-resistant lung adenocarcinoma cell lines

Next, we searched for information about copy number alteration or mutation of the 77 genes implicated in the abovemetioned altered biological functions in the crizotinib resistant cells. Deletions and mutations were present at a very low frequency. However, 26 genes were found to be amplified (> 1.5%), suggesting a high potential role for these genes as therapeutic targets in LUAD (Table 2).

**Table 2.** List of amplified genes. List of 26 genes with copy number amplification (>1.5%) among 77 genes upregulated in resistant cell lines. The table reports the frequency of mutation and deletion (percentage and number of cases in brackets) and known functional annotations. Functional categories: 1. Post-translational protein modifications, 2. Regulation of spliceosome assembly, 3. Regulation of Golgi transport, vesicle recycling and protein localization, 4. Regulation of transcriptional and translational processes, 5. Modulation of energy supplied by mitochondria and apoptotic processes, 6. Regulation of metabolic pathways, 7. Regulation of autophagy, 8. Negative regulation of adipogenesis.

| Pan-Lung Cancer (TCGA, Nat Genet 2016) n=1144. Lung Adenocarcinoma |  |  |  |  |  |  |  |  |  |  |  |
| --- | --- | --- | --- | --- | --- | --- | --- | --- | --- | --- | --- |
| Genes | Amplification | Mutation | Deletion | Function |  |  |  |  |  |  |  |
|  |  |  |  | 1 | 2 | 3 | 4 | 5 | 6 | 7 | 8 |
| AKT2 | 1.97%(13) | 1.06% (7) | - | X |  |  |  | X | X |  |  |
| DYRK1B | 1.67%(11) | 0.76% (5) | - | X |  |  |  |  |  |  |  |
| STX16 | 3.64%(24) | 1.06% (7) | - | X |  |  |  |  |  |  |  |
| RAB22A | 3.48%(23) | - | - | X |  | X |  |  |  |  |  |
| TM9SF4 | 1.52%(10) | 1.06% (7) | - | X |  |  |  |  |  |  |  |
| PRPF6 | 4.85%(32) | - | 0.15% (1) |  | X |  |  |  |  |  |  |
| SRSF6 | 1.52%(10) | 0.91% (6) | 0.15% (1) |  | X |  |  |  |  |  |  |
| ARFGEF2 | 1.82%(12) | 2.58% (17) | 0.15% (1) |  |  | X |  |  |  |  |  |
| CNIH3 | 4.24%(28) | 0.61% (4) | - |  |  | X |  |  |  |  |  |
| ADRM1 | 3.64%(24) | - | - |  |  |  | X |  |  |  |  |
| CSTF1 | 2.58%(17) | 1.21% (8) | - |  |  |  | X |  |  |  |  |
| NELFCD | 3.18%(21) | 0.76% (5) | - |  |  |  | X |  |  |  |  |
| TAF4 | 3.33%(22) | 1.97 (13) | - |  |  |  | X |  |  |  |  |
| TCFL5 | 3.94%(26) | 0.61% (4) | 0.15% (1) |  |  |  | X |  |  |  |  |
| YTHDF1 | 3.94%(26) | 1.67% (11) | 0.15% (1) |  |  |  | X |  | X |  |  |
| EIF3H | 4.24%(28) | 0.3% (2) | - |  |  |  | X |  |  |  |  |
| EIF3E | 3.94%(26) | 1.52% (10) | 0.3% (2) |  |  |  | X |  | X |  |  |
| RIDA | 4.7%(31) | 0.3% (2) | - |  |  |  | X |  |  |  |  |
| MTG2 | 3.64%(24) | 1.21% (8) | - |  |  |  | X |  |  |  |  |
| NDUFB9 | 6.82%(45) | 0.3% (2) | - |  |  |  |  | X |  |  |  |
| NDUFB5 | 2.12%(14) | 0.76% (5) | 0.45% (3) |  |  |  |  | X |  |  |  |
| SIRT2 | 2.27%(15) | 0.15% (1) | - |  |  |  |  |  | X | X | X |
| DPM1 | 1.82%(12) | 0.76% (5) | 0.15% (1) |  |  |  |  |  | X |  |  |
| FOXK2 | 1.97%(13) | 0.76% (5) | 0.15% (1) |  |  |  |  |  | X | X |  |
| TFEB | 2.88%(19) | 0.61% (4) | - |  |  |  |  |  |  | X |  |
| LRP3 | 3.03%(20) | 0.3% (2) | - |  |  |  |  |  |  |  | X |

Then, we intended to search for an association between these amplified genes and patient outcome. Using KM plotter online tool, that enclosed information for 1308 LUAD patients, we identified 9 genes significantly associated with detrimental patient outcome in terms of OS or PFS: DYRK1B, *TM9SF4, NDUFB9, SIRT2, CNIH3, ADRM1, NELFCD, MTG2*, and *TFEB* (Fig. 2 and 3).

**Figure 2.**
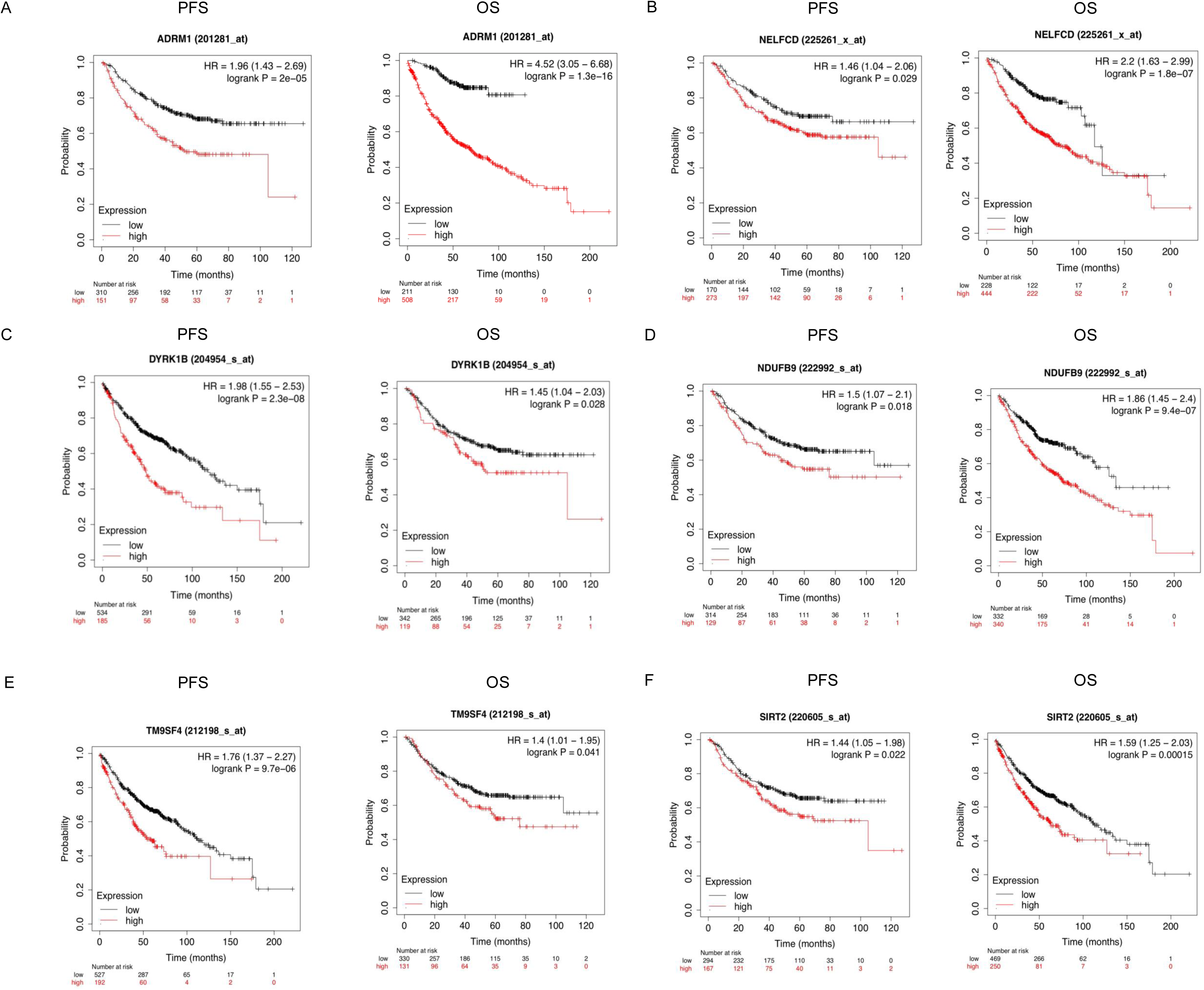
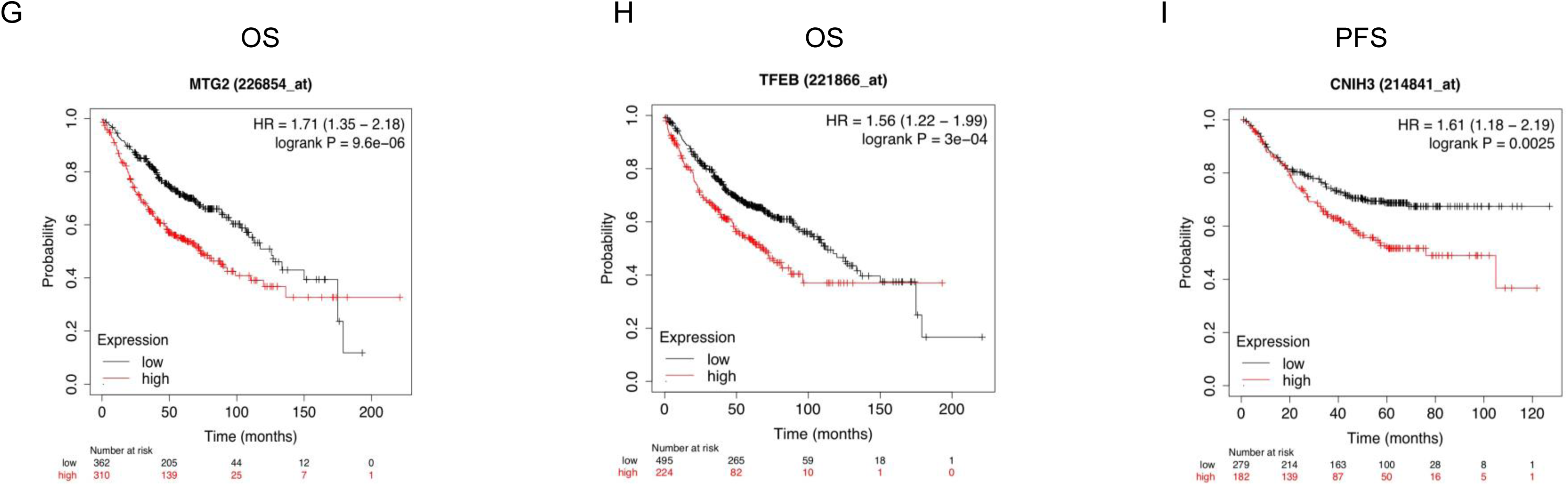
Association between gene expression and prognosis. Kaplan–Meier plots showing the association between gene expression and overall survival (OS) and progression-free survival (PFS). A. DYRK1B. B. TMPSF4. C. ADRM1. D. NELFCD. E. NDUFB9. F. SIRT2. Kaplan–Meier plots showing the association between gene expression and overall survival (OS) and progression-free survival (PFS). G. MTG2. H. TFEB. I. CNIH3

*ADRM1* amplification was associated with both worse OS (HR 4.52, 3.05-6.68, p value 1.3-16) and PFS (HR 1.96, 1.43-2.69, p value 0.00002) with an FDR (false Discovery rate of 1%) (Fig. 2A). *NELFCD* was associated with worse OS (HR 2.21, 1.63-2.99, p value 1.8-7 FDR 1%) and PFS (HR 1.46, 1.04-2.06, p value 0.029, FDR 50%) (Fig. 2B); *DYRK1B* was associated with worse OS (HR 1. 98, 1.55-2.53, p value 2.3-8, FDR 1%) and PFS (HR 1.45, 1.04-2.03, p value 0.028, FDR 50%). (Fig. 2C) *NDUFB9* was found to be associated with worse OS (HR 1.86, 1.45-2. 4, p value 9.4-7, FDR 1%) and PFS (HR 1.5, 1.07-2.1, p value 0.018, FDR 50%) (Fig. 2D). *TMPSF4* was related to worse OS (HR 1.76, 1.37-2.27, p value 9.7-6, FDR 1%) and PFS (HR 1. 41, 1.01-1.95, FDR 50%) (Fig. 2E). SIRT2 was related to worse OS (HR 1.59, 1.25-2.03, p value 0.00015, FDR 5%) and PFS (HR 1.44, 1.05-1.98, p value 0.022, FDR 50%) (Fig. 2F).

*MTG2* was found to be related to worse OS (HR 1.71, 1.35.2.18, p value 9.6-6, FDR 1%), but not to worse PFS (HR 0.86, 0.61-1.21, p value 0.39, FDR 100%) (Fig. 2G). *TFEB* is related to worse OS (HR 1.56, 1.22-1.99, p value 3-4, FDR 10%), but not to worse PFS (HR 0.76, 0.54-1.07, p value 0.11, FDR 100%) (Fig. 2H). CINH3 is associated with worse PFS (HR 1.61, 1.18-2.19, p value 0.0025, FDR 50%), but not with worse OS (HR 0.77, 0.61-0.97, p value 0.028, FDR 50%) (Fig. 2I).

### 3.3. Association of crizotinib resistant genes with immune infiltration

In cancer biology, not only the tumor cell plays a role, but also the TME where we find immune cells or angiogenesis processes, among others. Therefore, the knowledge of immune infiltration in the TME could help us to better understand the biology of these tumors and to propose therapeutic strategies that modulate this immunity. For instance, a high presence CD8 expressing lymphocytes is associated with increased immune processes of tumor recognition and death, and therefore with better prognosis [12]. In sharp contrast, tumor-associated macrophages (TAMs) mediate immunosuppressive effects on the adaptive immune cells of the TME, as TAMs display an ability to suppress T cell recruitment and function as well as to regulate other aspects of tumor immunity, what eventually results in supporting cancer cell growth and metastasis [13]. Moreover, T-regs are required to limit autoimmunity, so they can be deleterious in cancer through suppression of anti-tumor immunity [14]. Neutrophils are also related to a worse prognosis, because they stimulate a transition towards a mesenchymal phenotype characterized by an increase in cell migration and metastasis [15]. Finally, dendritic cells often regulate anti-tumor CD8 T cell immunity, leading to tolerance to this population and, therefore, limiting their function [16]. Cancer-associated fibroblasts (CAFs) are the principal component of stromal cells and release inflammatory, growth factors, and extracellular matrix, accelerating tumor proliferation and contributing to therapy resistance [17]

In our study, we found that three of the studies genes, TM9SF4, CNIH3, and ADRM1, positively correlate with at least two immunosuppressive populations while negatively correlate with CD8+ T-cells (Table 3). In short, *TM9SF4* had a positive correlation with neutrophils (Pearson correlation coefficient, PCC=0.182), fibroblasts (PCC=0.298) and negative correlation with CD8 (PCC=-0.133). *CNIH3* displayed a positive correlation with neutrophils (PCC=0.229), macrophages (PCC=0.256), fibroblasts (0.323), and DC (PCC=.349), and a negative correlation with CD8 (PCC=-0.199). *ADRM1* showed a positive correlation with T-Regs (PCC=0.199) and fibroblasts (PCC=0.131) and negative correlation with CD8 (PCC=-0.148). Although no negative correlation with the CD8+ T cell population was found, SIRT2 and TFEB expression was also associated with a higher infiltration of immunosuppressive cells (Table 3). Thus, *SIRT2* showed a positive correlation with neutrophils (PCC=0.112), macrophages (PCC=0.233) and DC (PCC=0.204). *TFEB* had a positive correlation with T-regs (PCC=0.235), macrophages (PCC=0.163), DC (PCC=0.262) and CD8 (PCC=0.124). No clear association with either immunosuppressive or immunoreactive populations was found for the rest of the genes (Table 3). *DYRK1B* just showed a positive correlation with T-regs immune infiltration (PCC =0.18) and a non-significant positive correlation with fibroblasts (PCC=0.05). *MTG2* appeared to have a positive correlation with T-regs (PCC=0.104), a negative correlation with macrophages (PCC=-0.105), and a non-significant positive correlation (PCC=0.012) with fibroblasts. *NELFCD* had a negative correlation with neutrophils (PCC=-0.102), macrophages (PCC= −0.205), and DC (PCC=-0.247) and a non-significant positive correlation with fibroblasts (PCC=0.072). Last, NDUFBP had a negative correlation with neutrophils (PCC=-0.247), fibroblasts (PCC=-0.119) and DC (PCC=-0.262) (Table 3).

**Table 3.** Association between gene expression and immune cell infiltration. Positive correlations (Spearman’s r > 0, p < 0.05) are highlighted in green; negative correlations (Spearman’s r < 0, p < 0.05) in red. Non-significant associations (p > 0.05) are uncolored. Data primarily obtained from TIMER 2.0 using EPIC expression profiles; results based on TIMER are indicated by a dash (-). T-regulatory cell infiltration was analyzed with CIBERSORT.

| Gene | T-Regs | Neutrophils | Macrophages | Dendritic cells | T-Cell CD8 |
| --- | --- | --- | --- | --- | --- |
| *DYRK1B* | 0.18 | -<br>-0.085 | -0.073 | -0.055 | 0.081 |
| *TM9SF4* | 0.05 | -<br>0.182 | -0.09 | -0.01 | -0.133 |
| *CNIH3* | 0.1 | -<br>0.229 | 0.256 | -<br>0.349 | -0.199 |
| *ADRM1* | 0.199 | -<br>-0.058 | -0.044 | -<br>-0.012 | -0.148 |
| *NELFCD*<br>(*TH1L*) | 0.032 | -<br>-0.102 | -0.205 | -<br>-0.247 | -0.053 |
| *MTG2*<br>(*GTPBP5*) | 0.104 | -<br>-0.051 | -0.105 | -<br>-0.091 | -0.063 |
| *NDUFB9* | 0.013 | -<br>-0.247 | -0.097 | -0.262 | -0.075 |
| *SIRT2* | 0.077 | -<br>0.112 | 0.233 | 0.204 | -0.011 |
| *TFEB* | 0.235 | -<br>0.035 | 0.163 | -<br>0.262 | 0.124 |

### 3.4. Identification of potential drugs for the modulation of these genes

Finally, having identified at least five genes that are overexpressed in crizotinib-resistant cell lines, associated with worse prognosis and with an immunosuppressive TME, we have sought treatments that inhibit their expression, with the hypothesis that this could ultimately improve the survival of LUAD patients who display resistance to this ALK inhibitor alone or in combination with immunotherapy (Table 4). *ADRM1*-inhibiting molecules include bortezomib, approved for the treatment of multiple myeloma, carfilzomib, oprozomib, and RA-190 [18]. *SIRT2* inhibitor molecules include gefitinib, approved for EGFR-mutated lung cancer, and veliparib. Other molecules under investigation for *SIRT2* targeting are AEM1 and AEM2 [19,20,21]. There is not much therapeutic evidence for the inhition of other genes, such as CNIH3 and TM9SF4, with only opioids [22] and miR-1226-3p [23], as potential targeting drugs, respectively.

**Table 4.** Candidate therapeutic targets and availability status. Availability status of the therapeutic agents (approved for other indications or investigational) against the nine upregulated genes identified as potential therapeutic targets.

| Therapeutic agents |  |  | Pubmed references (lung cancer and gene) |
| --- | --- | --- | --- |
| Genes | Approved for other indications | In research |  |
| DYRK1B | Fostamatinib, Palbociclib | Harmine, Cenisertib, Voruciclib, AZ191, SM08502 | PMID: 25982285<br>PMID: 23311607<br>PMID: 19633423<br>PMID: 20512148<br>PMID: 25963666 |
| TM9SF4 |  | miR-1226-3p |  |
| NDUFB9 | NADH, FAD, FE<sup>+2</sup>, NAD, ubiquinone-1 |  |  |
| SIRT2 | Gefitinib, nicotinamide | Cambinol, NAD, resveratrol, veliparib, SirReal2, tenovin 6, selisistat, tenovin 3, AGK2, thiomyristoyl, salermidem, SRT 1720 HCl, OSS 128167, AK7, sirtinol, 3-TYP, tenovin 1, AEM1, AEM2 | 20 references |
| CNIH3 | Opioids |  |  |
| ADRM1 | Bortezomib, carfilzomib, | Oprozomib, RA-190 |  |
| NELFCD | Hidrochlorotiazide |  |  |
| MTG2 |  | Guanosine triphosphate, |  |
| TFEB |  |  | PMID: 30013069<br>PMID: 26264650 |

Although, no association with and immunosupressive TME was found for *DYRK1B* overexpression, still, targeting this gene could be of interest in the clinical setting, as this gene had the higher association with bad prognosis in LUAD. In fact, there are agents approved for indications other than lung cancer, such as fostamatinib or palbociclib, that can target this gene. Other agents such as harmine, cenisertib, or voruciclib are under study [24,25,26].

### 3.5. Differential expression between normal and tumoral tissue

To complete our study, we investigated the potential differential expression between normal and tumoral tissue for the 26 overexpressed genes that showed an amplification >1.5% using TNM plot, as a treatment focused on a gene that is overexpressed in tumoral tissue would have a better toxicity profile with less interaction with normal tissue. Our analysis revealed that most of the identified genes are overexpressed in tumoral tissue, except of STX6, *SIRT2, TFEB,* and AKT2 (Table 5), which are overexpressed in normal tissue.

**Table 5.**
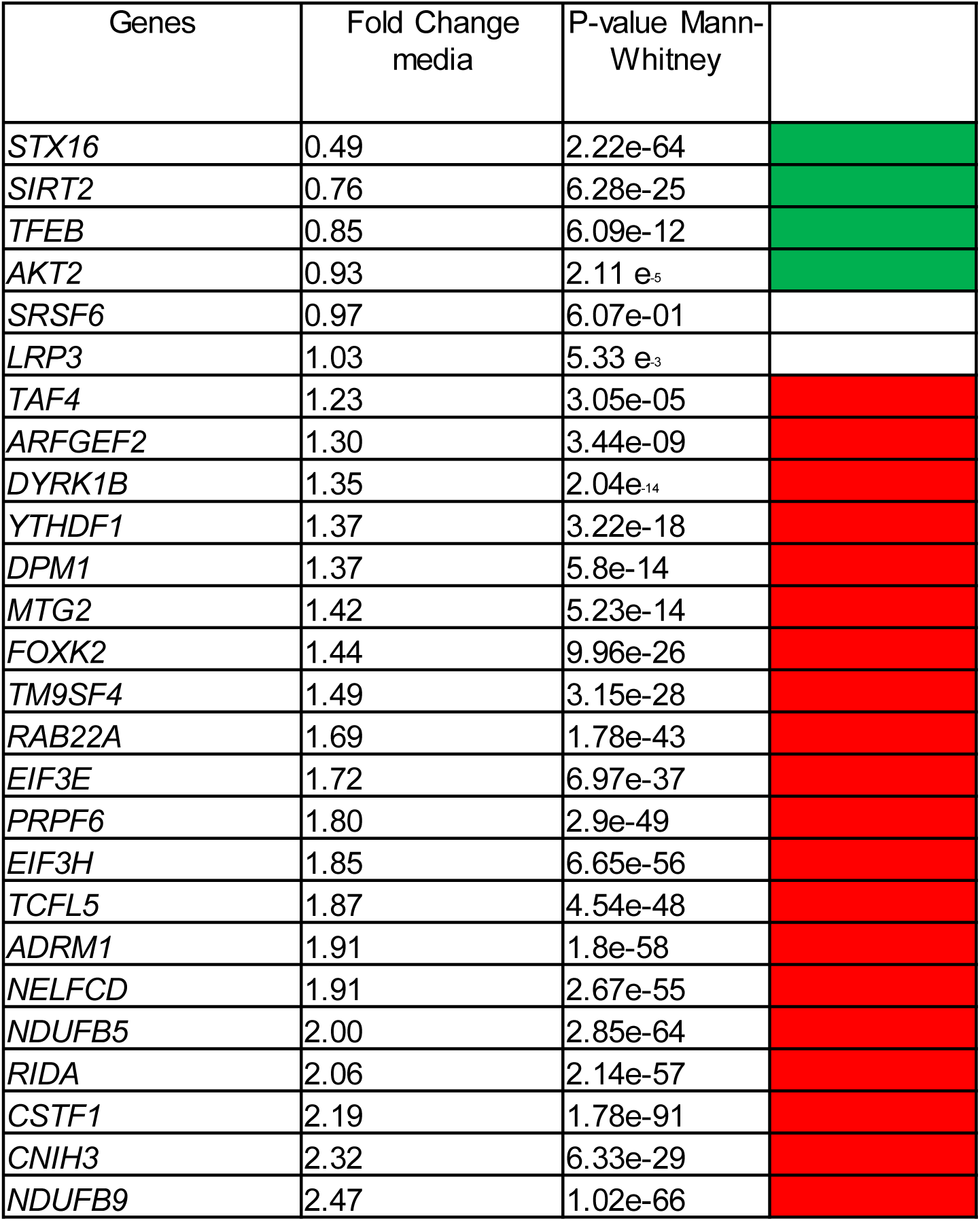
Differential expression of amplified genes in LUAD. . Differential expression of the 26 amplified genes (>1.5%) between non-tumoral and LUAD tumor tissues. Fold-change (FC) values are shown: FC > 1 indicates higher expression in tumor tissue (red); FC < 1 indicates higher expression in normal tissue (green). Non-significant results are uncolored. Genes are ordered by fold-change magnitude.

## 4. Discussion

In this study, we identified nine amplified genes in crizotinib-resistant LUAD cell lines, whose expression is associated with poor prognosis: *DYRK1B, TM9SF4, NDUFB9, SIRT2, CNIH3, ADRM1, NELFCD, MTG2,* and *TFEB*. Our findings suggest that these genes are involved in critical cell cycle functions and are amplified in over 10% of LUAD, with some cases showing amplification rates as high as 45%. This highlights the potential relevance of targeting these genes with inhibitors. Additionally, we observed a correlation between the expression of five of these genes, *SIRT2, TFEB, CNIH3, ADRM1,* and *TM9SF4* and infiltration of immune population, suggesting that patients with overexpression of these biomarkers could benefit from combinational treatment of crizotinib and immunotherapy or drugs targeting these genes. could be a promising treatment strategy for crizotinib-resistant tumors overexpressing these five genes, suggesting its use as potential biomarkers in LUAD.

Gene amplification results in the duplication of chromosomal fragments of different sizes, which could lead to a potential advantage in tumor cell growth and an important mechanism of resistance to antitumor treatments. Therefore, the identification of amplified genes associated with poor disease prognosis in LUAD could be useful to uncover novel potential targets which might results in an improvement of therapy. In this study we have observed that elevated RNA expression of six of out the nine amplified genes (*DYRK1B, TM9SF4, ADRM1, NELFCD, NDUFB9, SIRT2*) would imply a negative prognostic impact in both PFS and OS. Conversely, *MTG2* and *TFEB* are associated with poor prognosis only in OS and *CNIH3* only with PFS.

Knowing how the TME is organized and which immune population are the most represented could be useful to better understand the biology of tumors and, hence, to propose potential therapeutic strategies that modulate their immunity. For instance, the overexpression of CD8 in TME enhances the immunogenicity of tumors, whereas the overexpression of DC, T-regs, macrophages, and neutrofils are associated with an immunosuppressive TME. Indeed, in metastatic non-small cell lung cancer, there are data supporting that a higher infiltration by CD8 cells correlates with better OS, as opposed to a high infiltration by T reg lymphocytes [27]. Moreover, the presence of tumor-infiltrating CD8 lymphocytes has been associated with greater response to checkpoint inhibitors and, therefore, with better patient prognosis [28]. In this scenario, we hypothesized that tumors with a suppressive TME could be less sensitive to immunotherapy in monotherapy, but this immune response could be enhanced using a combination treatment of immunotherapy with other agents, such as anti-angiogenic (bevacizumab, aflibercept) or classical chemotherapy (platinum compounds). Then, the association between immune infiltrates and gene expression was investigated (Table 3).

In our study, we found that three of the investigated genes, TM9SF4, CNIH3, and ADRM1, positively correlate with at least two immunosuppressive populations while negatively correlate with CD8+ T-cells (Table 3). In addition, SIRT2 and TFEB expression was also associated with a higher infiltration of immunosuppressive population, each of them correlating with three immunosuppressive cell types (Table 3). Then, tumors with increase expression of one or more of these five markers could benefit from combination therapies with immunotherapy and other treatments, such as antiangiogenic o classic cytotoxic, or even available drugs targeting these biomarkers (Table 4).

TMPFS4 is a gene that encodes for a transmembrane protein related to protein localization processes, regulation of intracellular pH, or vesicular transport (Figure 1) [29]. This gene, which is overexpressed in melanoma, colon cancer, gastrointestinal tumors, acute myeloid leukemia, myelodysplastic syndrome, and breast cancer, is associated with resistance to anthracycline regimens in some contexts [29,30]. *CNIH3*, which encodes a transmembrane protein related to transport (Figure 1) [31], is overexpressed in melanomas, gliomas, and squamous esophageal cancer [31]. In the latter localization, preliminary attempts have been made to demonstrate its potential in discerning between healthy tissue and tumor tissue [31]. Last, *ADRM1*, which encodes a protein related to cell adhesion processes (Figure 1) [18], is overexpressed in different solid tumors such as hepatocarcinoma, intrahepatic cholangiocarcinoma, gastric cancer, luminal breast cancer, or ovarian cancer [18,32,33]. Among the molecules that can potentially inhibit *ADRM1* is bortezomib (Table 4), approved for the treatment of multiple myeloma. SIRT2 is overexpressed in breast, liver, colon, glioma, renal, gastric, melanoma, ovarian or prostate cancer [19,20]. This gene encodes for a protein deacetylase that catalyzes the elimination of the acetyl group of numerous proteins and participates in regulation of metabolic pathways, autophagy and adipogenesis (Figure 1) [19], and whose action at the level of lung tumor cells is ambiguous, because while some studies points at it as a suppressor gene, others suggest it acts as an oncogene [19,20]. In fact, inhibition of SIRT2 could have a protective effect in patients with high SIRT2 expression and increase sensitivity to etoposide [19]. SIRT2 inhibitor molecules include gefitinib, approved for EGFR-mutated lung cancer, and veliparib (Table 4). Other molecules under investigation are AEM1 and AEM2 (Table 4) Last, TFEB promotes DNA transcription and participates in different processes such as lysosomal localization or autophagy (Figure 1). In the case of KRAS and LKB1 mutated lung cancers, inhibition of acetylation via TFEB reduces metastatic capacity *in vivo* [34].

Regarding the rest of the nine genes initially identified in this work, *DYRK1B* has demonstrated its pro-tumor effect in cellular models of the ovary, pancreas, glioblastoma, colon, melanoma, or breast [24]. Its expression is stimulated in stressful situations, such as chemotherapy exposure, favoring tumor cells to enter quiescence [24, 25]. Up to 90% of lung tumors overexpress this protein, and its depletion increases cell sensitivity to cisplatin-induced apoptosis [26]. At the therapeutic level, there are agents approved for indications other than lung cancer, such as fostamatinib or palbociclib. Other agents such as harmine, cenisertib or voruciclib are under study (Table 4). *NDUFB9*, is associated with mitochondrial oxidative metabolism (Figure 1) and is found to be amplified in other tumors such as breast cancer, uveal and cutaneous melanoma [35]. Metformin and Fe +2 have been shown to interfere with *NDUFB9* expression (Table 4). *MTG2*, which acts in ribosomal assembly (Figure 1), is overexpressed in malignant gliomas, breast, prostate, kidney, pancreas, colorectal and liver cancers [36]. Last, the NELFCD complex inhibits transcriptional elongation of RNA polymerase II (Figure 1) [37] and is overexpressed in colorectal cancer. endometrial, liver or prostate cancer [37]. Although some of these candidates could have potential as therapeutic targets in LUAD, no clear association with either immunosuppressive or immunoreactive populations was found in this study (Table 3) and, therefore, patients with overexpression of these four genes might not benefit from combinational therapy with immune drugs

Finally, therapeutic strategies development must take into consideration the toxicity profile, to avoid side effect on normal tissues. Then, treatments focused on genes overexpressed in tumor tissues in comparison with non-pathological ones would have a better toxicity profile than highly expressed in normal tissue (Table 5). However, two of our candidates, *SIRT2,* and *TFEB,* were found to be overexpressed in normal tissue. For this reason, with the aim of adjusting doses and detecting possible side effects as soon as possible, more exhaustive studies of pharmacodynamics and pharmacovigilance of treatments aimed at these targets should be performed in the future to confirm them as potential targets in LUAD.

### 4.1. Conclusion

In this study, we identified five genes, *SIRT2, TFEB, CNIH3, ADRM1,* and *TM9SF4*, associated with resistance to crizotinib treatment, poor prognosis in LUAD and potential sensitivity to immunotherapy, due to their association with infiltration of immunosuppressive populations. Therefore, combining crizotinib with immunotherapy or targeted therapies with drugs against the molecular pathways in which this biomarkers are involved may offer improved therapeutic outcomes for patients with tumors expressing these genes.

The identification of these genes also opens new possibilities for combining crizotinib with drugs targeting these molecular pathways, some of which are already approved in other contexts, including lung cancer (e.g., gefitinib). Additionally, their overexpression has significant implications for the TME, which could guide the development of therapeutic strategies incorporating immunomodulatory treatments.

Although our findings were derived from *in silico* analysis, with the inherent limitations of such approach, they represent a promising avenue for future research. These results could be further validated and translated into clinical practice to improve treatment outcomes in LUAD.

Our study identified nine genes that are overexpressed in crizotinib-resistant ALK-mutated lung adenocarcinoma cell lines and are linked to worse prognosis. Additionally, we found significant variations in immune infiltration patterns in the tumor microenvironment, depending on the overexpressed gene. These findings provide valuable insights into the mechanisms of crizotinib resistance and suggest potential strategies for overcoming this resistance. Combining crizotinib with immunotherapy or targeted therapies against specific resistance pathways may offer improved therapeutic outcomes for patients with tumors expressing these genes.

## Declaration of competing interest

The authors declare that they have no know competing financial interests/personal relationships which may be considered as potential competing interests.

## Author contribution

**Édgar Villena:** Conceptualization, Data curation, Formal analysis, Investigation, Methodology, Validation, Visualization, Writing - original draft, and Writing - review & editing. **Maissaa Bouchakour** Visualization. P**aula Sánchez Olivares:** Visualization, Writing - review & editing. **Fernando Andres-Pretel:** Data curation; Software. **Eva M. Galán-Moya:** Conceptualization, Data curation, Formal analysis, Investigation, Methodology, Project administration, Resources, Software, Supervision, Validation, Visualization, Writing - original draft, and Writing - review & editing.

## Acknowledgements

We thank all the members of the Cancer Pathophysiology & Therapy Lab for their support.

## Abbreviations

*ALK*: Anaplastic Lymphoma Kinase
EMA: European Medicines Agency
EML4: echinoderm microtubule-like protein 4
FDA: Food and Drug Administration
IC50: half maximal inhibitory concentracion
KIF5B: Kinesin Family Member 5B
JAK-STAT: Janus Kinasa
KLC1: Kinesin Light Chain 1
KMp: Kaplan Meier plotter
LUAD: lung adenocarcinoma
MAPK: Mitogen-Activated Protein Kinase
PI3K: Phosphoinositide 3-kinases
PFS: progression free survival
OS: overall survival
TCGA: Cancer Genome Atlas
TKI: tyrosine kinase inhibitor
TGF: tumor growth factor.

## Supplementary material

**Supplementary Table 1.**
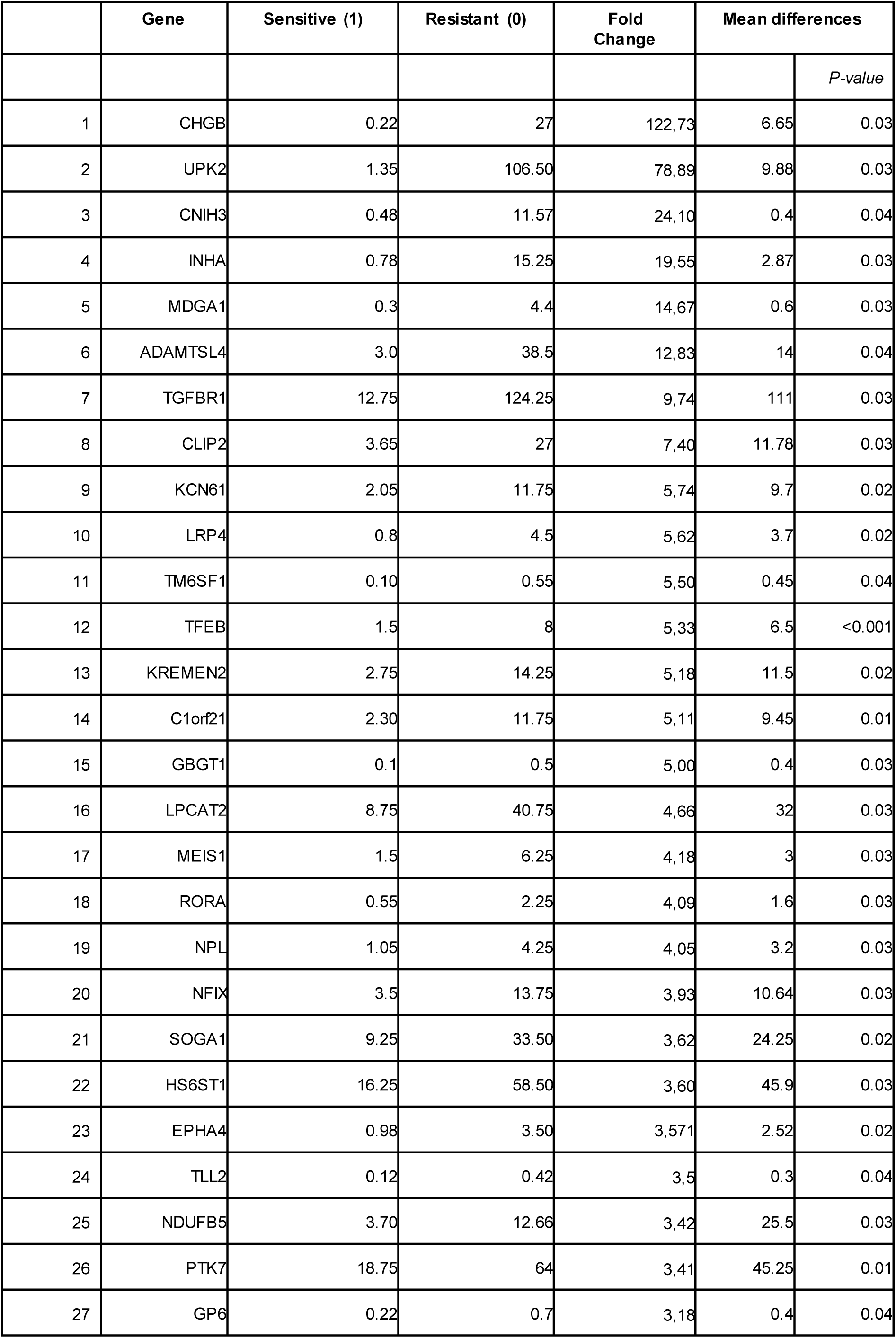
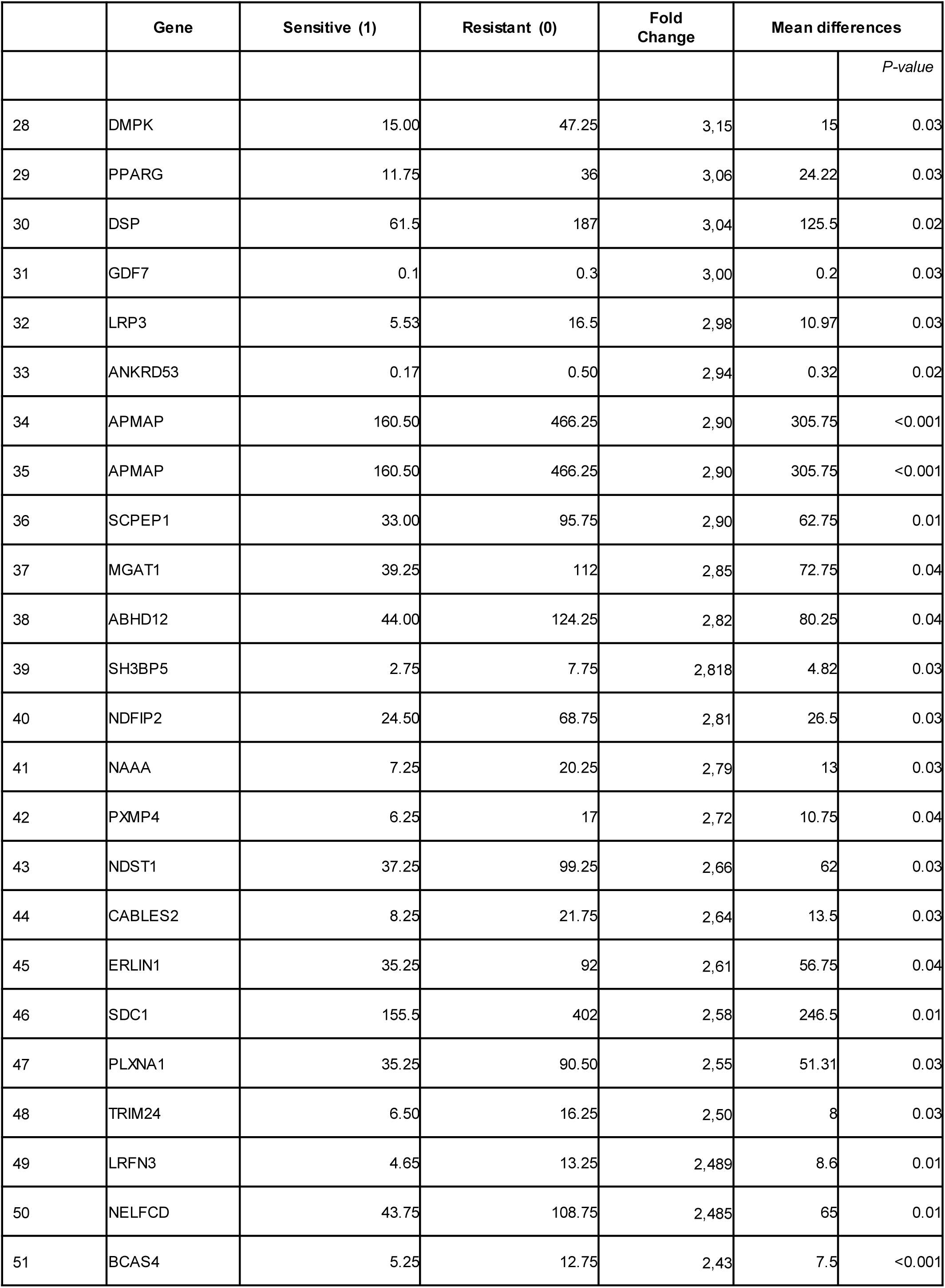
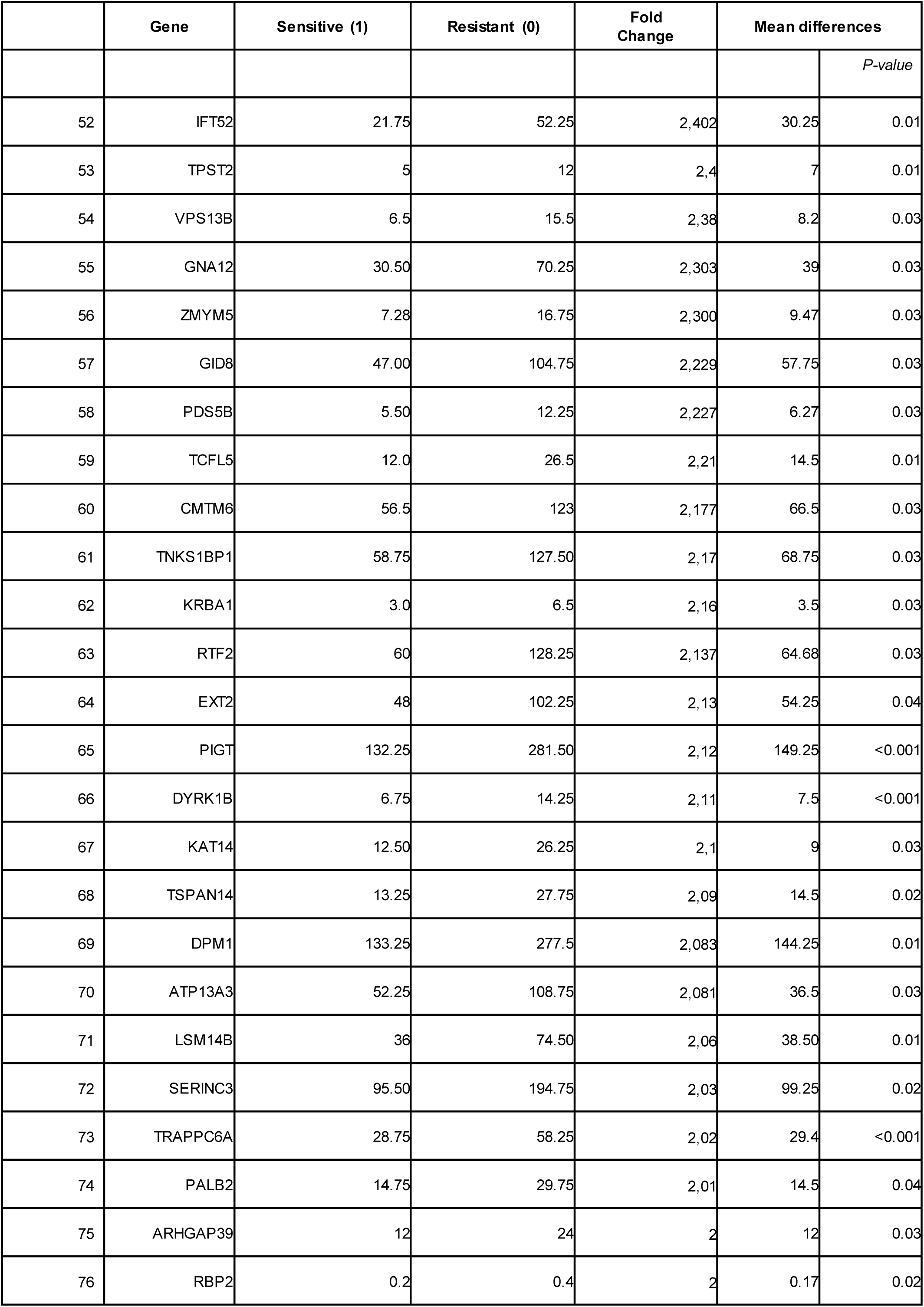
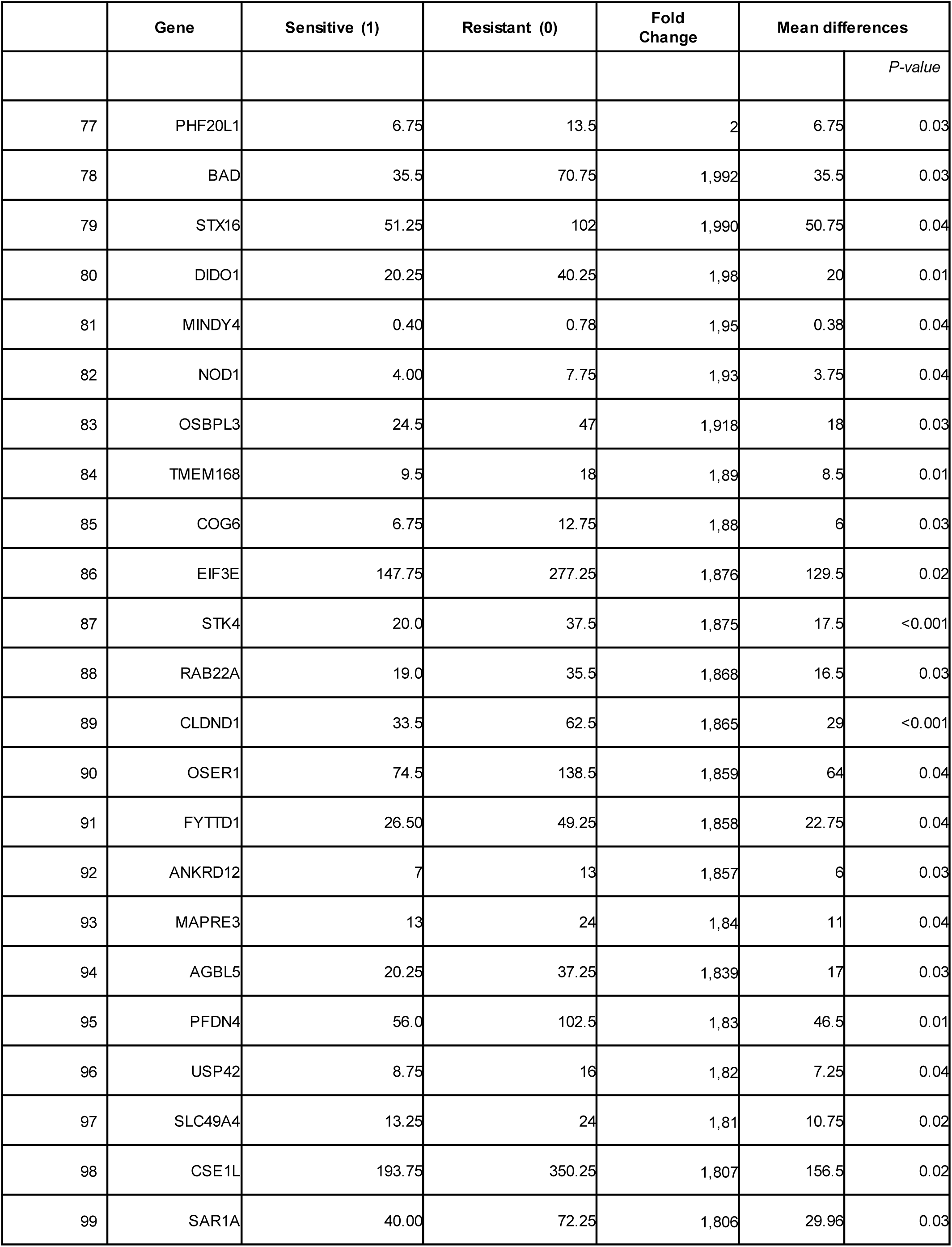
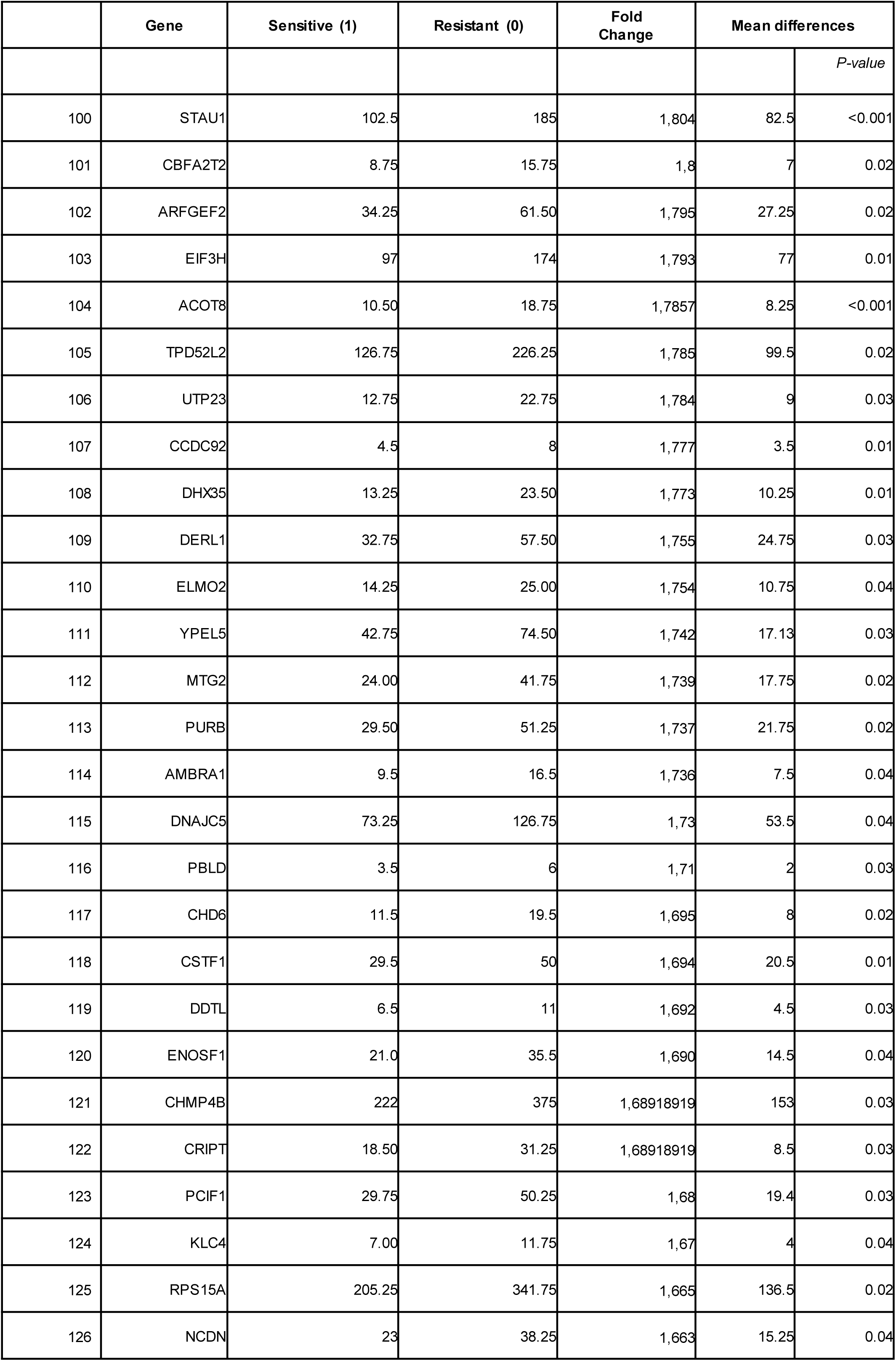
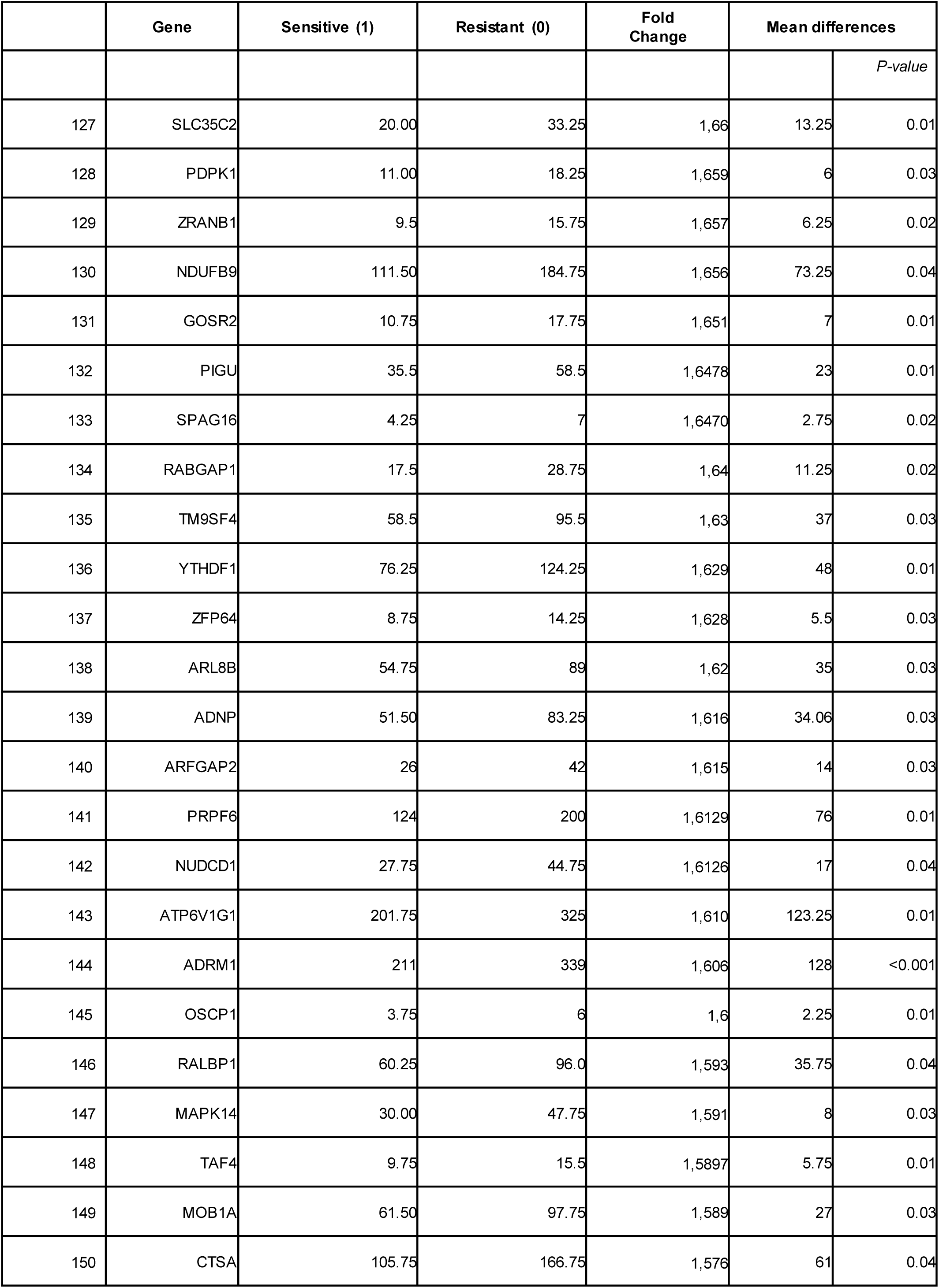
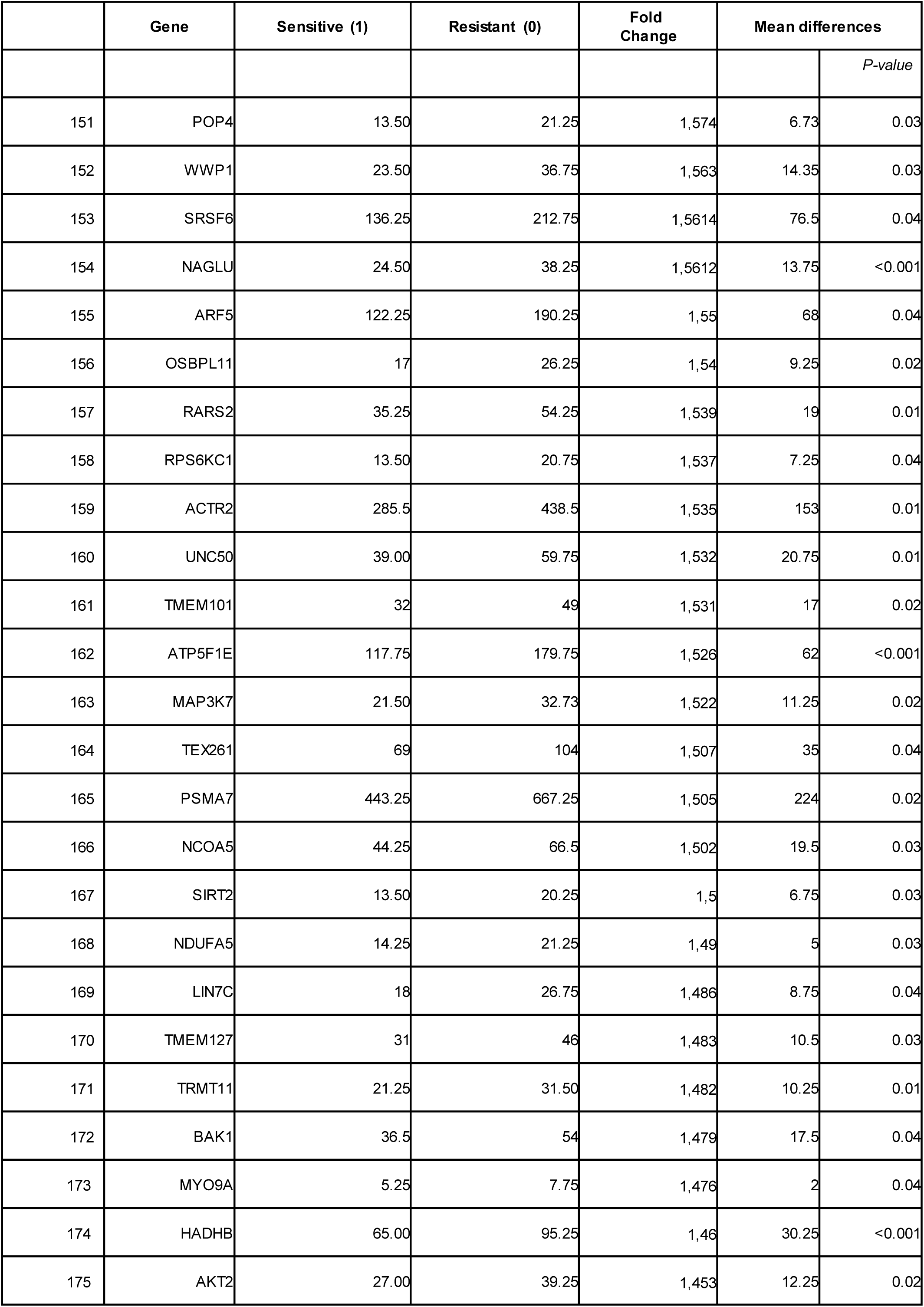
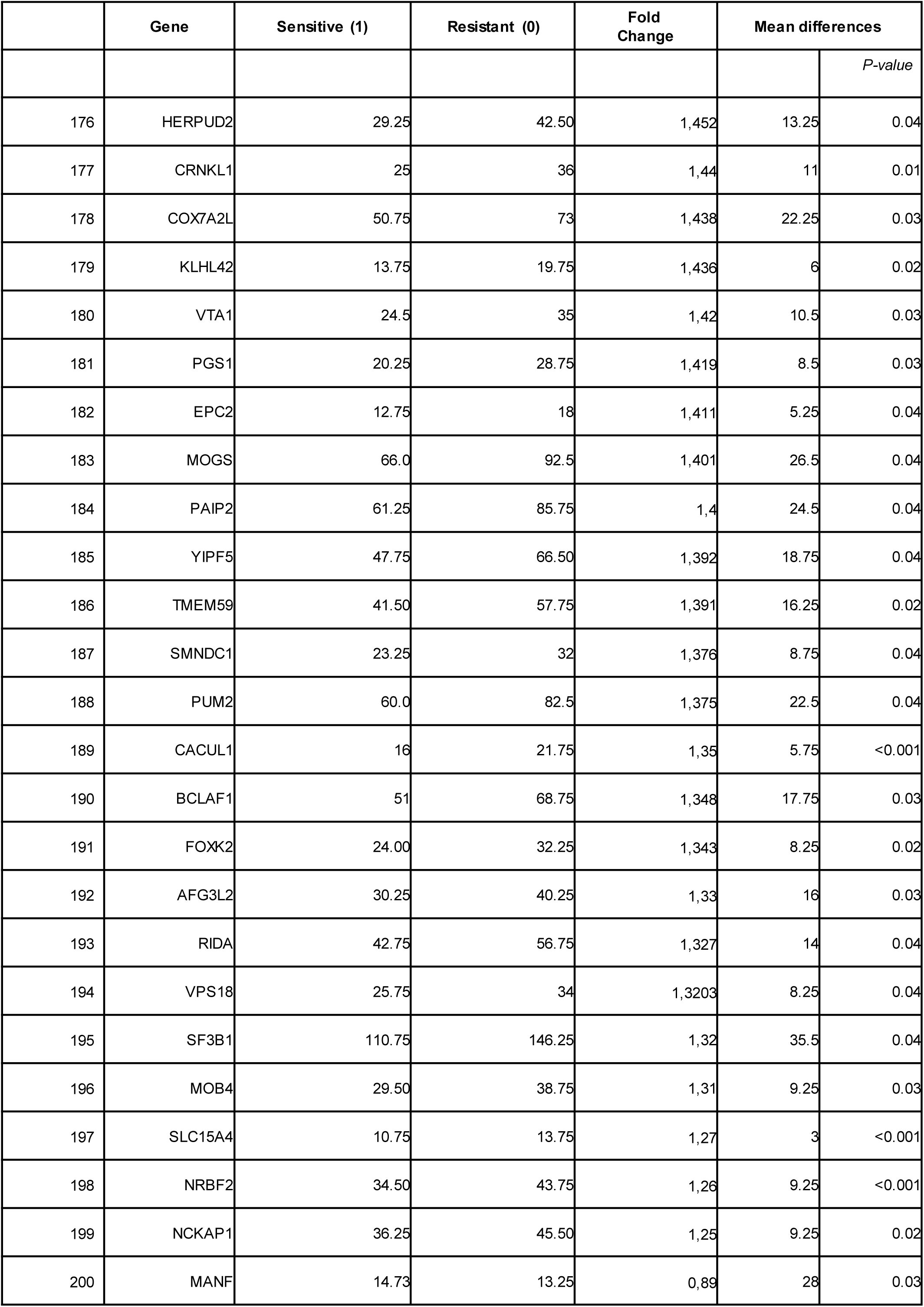
Gene expression differences between sensitive and resistant lung adenocarcinoma cell lines. Values are expressed as the median difference in expression between both groups. Genes are ordered by fold change, from highest to lowest. Only genes with statistically significant differential expression (p ≤ 0.05) are included.

## Notes

### Competing Interest Statement

The authors have declared no competing interest.

